# Centromere-proximal meiotic crossovers in *Drosophila melanogaster* are suppressed by both highly-repetitive heterochromatin and the centromere effect

**DOI:** 10.1101/637561

**Authors:** Michaelyn Hartmann, James Umbanhowar, Jeff Sekelsky

## Abstract

Crossovers are essential in meiosis of most organisms to ensure the proper segregation of chromosomes. The lack or improper placement of crossovers can result in nondisjunction and aneuploidy in progeny. Crossovers near the centromere can cause nondisjunction; centromere-proximal crossovers are suppressed by what is termed the centromere effect, but the mechanism is unknown. Here, we investigate contributions to centromere-proximal crossover suppression in *Drosophila melanogaster*. We mapped a large number of centromere-proximal crossovers and find that crossovers are essentially absent from the highly-repetitive (HR)-heterochromatin surrounding the centromere but occur at a low frequency within the less-repetitive (LR)-heterochromatic region and adjacent euchromatin. Previous research suggested that flies that lack the Bloom syndrome helicase (Blm) lose meiotic of crossover patterning, including the centromere effect. Mapping of centromere-proximal crossovers in *Blm* mutants reveals that the suppression within the HR-heterochromatin is intact, but the centromere effect is lost. We conclude that centromere-proximal crossovers are suppressed by two separable mechanisms: the HR-heterochromatin effect, which completely suppresses crossovers in the HR-heterochromatin, and the centromere effect, which suppresses crossovers with a dissipating effect with distance from the centromere.

## Introduction

Crossovers are essential for the proper segregation of homologous chromosomes in meiosis, which is evidenced by the fact that chromosomes lacking a crossover commonly segregate improperly in meiosis I (Koehler, Boulton, et al. 1996). However, it is not only the presence of crossovers that is important, but also their proper placement along the chromosome. Koehler et al. also showed that apparent meiosis II nondisjunction events occurred primarily in oocytes that experienced a centromere-proximal crossover (Koehler, Boulton, et al. 1996). Similarly, cases of human trisomy 21 that appear to have arisen from meiosis II nondisjunction are associated with an increase in centromere-proximal crossovers (Koehler, Hawley, et al. 1996; Lamb et al. 1996). It has long been known that crossovers near the centromere are reduced in many organisms; this has been referred to as the centromere effect (or the spindle-fibre effect before centromeres were defined) (Beadle 1932).

Meiotic recombination is initiated by DNA double-strand breaks (DSBs), each of which can be repaired to give crossovers or noncrossover products through a tightly-controlled decision (Lake and Hawley 2016; Miller et al. 2016). In addition to the centromere effect, interference and assurance also govern crossover patterning. Interference is the phenomenon where one crossover suppresses the occurrence of another crossover in nearby (A. H. Sturtevant 1913; reviewed in Berchowitz and Copenhaver 2010). Assurance is the phenomenon in which each pair of homologous chromosomes almost always receive at least one crossover regardless of size (Wang et al. 2015; Mather 1937). The effect of these crossover patterning phenomena on DSB repair results in the typical crossover distribution where most crossovers occur in the middle to distal end of the chromosome and are decreased near the centromere. The mechanisms of these phenomena are largely undescribed and remain elusive. In this study, we use *Drosophila melanogaster* to gain insight into the centromere effect and how crossovers are suppressed in the centromere-proximal regions.

Approximately one third of each *Drosophila* chromosome is composed of highly-repetitive, peri-centromeric satellite sequence arrays that are heterochromatinized. There have been studies that suggest heterochromatin plays a role in decreasing crossovers in the pericentric regions. It has been proposed that there are no crossovers within heterochromatin simply due to the tightly packed chromatin not being accessible to proteins that either make DSBs or repair them into crossovers. Support for this comes from cytological studies where Mehrotra and McKim (2006) observed no DSBs colocalizing with the heterochromatic mark HP1. Additionally, dominant suppressor of position-effect variegation, *Su(var)* mutations, that likely cause heterochromatin to assume a more open structure, allow an increase in crossovers within the pericentromeric heterochromatin (Westphal and Reuter 2002). These results support the idea that suppression of crossovers near the centromere is due to exclusion of DSBs in heterochromatin.

However, early studies on the centromere effect involving chromosome rearrangements in *Drosophila* show that centromere-proximal crossover suppression extends beyond heterochromatin into the euchromatin. Mather (1939) showed that a euchromatic region moved closer to the centromere, but nearer to a smaller amount of heterochromatin, experienced a greater decrease in crossovers than did a region moved slightly farther away from the centromere, but nearer to a larger amount of heterochromatin. He suggested that the decrease in crossovers was due to proximity to the centromere rather than the proximity to heterochromatin. Yamamoto and Miklos (1978) studied *X* chromosomes in *Drosophila* that had large deletions of the pericentromeric heterochromatin, and showed that the larger the deletion, the farther the decrease in crossovers spread into the euchromatin. They concluded that centromere-proximal crossover suppression does not depend on the amount of heterochromatin, but on distance from the centromere. Nonetheless, the question still remains whether heterochromatin has the ability to decrease crossovers in adjacent euchromatic regions; we address that question in this work.

Heterochromatin is not homogeneous and may not behave uniformly throughout. In polytene chromosome spreads, heterochromatin has two distinct appearances that have been described: alpha-heterochromatin is the small, densely staining region of the chromocenter that is highly underreplicated in this tissue, whereas beta-heterochromatin is more diffusely staining and is moderately replicated (Gall 1973; Ashburner 1980; Laird, Hammond, and Lamb 1987; Miklos and Cotsell 1990). Heterochromatin is not homogeneous based on sequence composition. Regions of pericentric heterochromatin adjacent to the euchromatin are composed of blocks of transposable elements (TEs) with varying amounts of repeats and interspersed unique sequence. This has made it possible to assemble these regions in the reference genome (Hoskins et al. 2015). Chromatin domains identified in cell lines show that much of this sequence is heterochromatic or transcriptionally silent (Filion et al. 2010; Thurmond et al. 2019). In contrast, sequences closer to the centromere are highly repetitive, consisting largely of blocks of tandemly-arrayed satellite sequences. These have not been assembled to the reference genome, but long-read sequencing has permitted assembly of some satellite arrays (Khost, Eickbush, and Larracuente 2017). We will refer to the two types of heterochromatin as highly-repetitive (HR)-heterochromatin and less-repetitive (LR)-heterochromatin.

In this study, we investigate the role of the two types of heterochromatin and the centromere effect in suppressing pericentromeric crossovers. We show that centromere-proximal crossover suppression is mediated by both a (HR)-heterochromatin effect and the centromere effect. The HR-heterochromatin effect is restricted to the highly-repetitive heterochromatin, which presumably does not allow double strand breaks to occur and therefore, no crossovers can be formed in these regions. This study allows some insight into chromosome characteristics that could be contributing factors to the centromere effect and supports the idea that the centromere effect is a protein mediated meiotic mechanism.

## Materials and Methods

### Drosophila stocks

Flies were maintained on standard medium at 25°C. Mutant alleles that have been previously described include *Blm*^*N1*^and *Blm*^*D2*^(McVey et al. 2007). *Blm*^*N1*^*/Blm*^*D2*^mutants experience maternal-effect lethality, which was overcome using the *UAS::GAL4* system with the *matα* driver as previously described (Kohl, Jones, and Sekelsky 2012).

### Phenotypic crossover distribution assay

Crossover distribution on chromosome *X* was scored by crossing *y sc cv v g f* • *y*^*+*^/ *M*{*3xP3-RFP.attP’*}*ZH-20C* to *y sc cv v g f* • *y*^*+*^ males, where • *y*^*+*^ is a duplication of *y*^*+*^ onto the right arm of the *X*. Crossover distribution on chromosome *2*L was scored by crossing virgin *net dpp*^*d-ho*^ *dp b pr cn / +* female flies to *net dpp*^*d-ho*^ *dp b pr cn* homozygous males. Crossover distribution on chromosome *2*R was scored by crossing *net dpp*^*d-ho*^ *dp b pr cn vg/ +* to *net dpp*^*d-ho*^ *dp b pr cn vg* homozygous males. Crossover distribution on chromosome *3* was scored by crossing virgin *ru h th st cu sr e ca / +* females to *ru h th st cu sr e ca* homozygous males. Crossovers between *px* and *sp* were scored by crossing virgin *px sp / +* to homozygous *px sp* males and scoring crossovers. Additionally, *px bw*^*D*^ *sp / +*, and *px sp / bw*^*D*^ were crossed to *px sp* homozygous males for scoring this interval in a *bw*^*D*^ background. Crossovers in *Blm* mutants were scored same way as chromosome *2*L in wild-type. Each cross was set up as a single experiment with at least 20 vials set up and flipped after three days. After three more days, parents were emptied from second round of vials. All progeny were scored for parental and recombinant phenotypes for five days from all vials. Crossover numbers in flies are shown as cM where cM = (number of crossovers / total number of flies) * 100. Fisher’s Exact Test was used to compare total COs to total number of flies. See Table S1 for phenotypic crossover distribution data.

### SNP/indel crossover mapping

Crossovers were finely mapped near the centromere using SNP/indels between isogenized strains. First, centromere-proximal crossovers were identified by phenotypic markers on the chromosome. For all chromosomes, crosses were set up between a wild-type chromosome and a chromosome with recessive markers; females heterozygous for these were collected and crossed to males homozygous for the recessive markers, and progeny were scored. Crossovers were collected between *f* and *y+* on the *X* chromosome, between *b* and *vg* on chromosome *2*, and between *h* and *e* on chromosome *3*. Illumina whole-genome sequencing was performed on each isogenized strain and genomes were assembled to the *Drosophila melanogaster* reference sequence, *Dm*6 (Hoskins et al. 2015), using BBMap (version 37.93, Bushnell 2014). SNPs and indels were called in comparison to the reference sequence using SAMtools mpileup (Sversion 1.7, Li et al. 2009; Li 2011), and then compared between strains using VCFtools (version 0.1.14, Danecek et al. 2011). Primers were designed to amplify only the wild-type chromosome so that each SNP/indel could be genotyped. See Table S2 for list of primers and locations. See table S3 and S4 for crossover distribution results from fine mapping for WT and *Blm*, respectively.

### Drosophila whole mount ovary immunofluorescence

About ten three- to five-day old virgins were kept in a vial with yeast paste overnight with a few males. Ovaries were dissected in phosphate buffered saline (PBS) and incubated for 20 minutes in fixative buffer (165 µL fresh PBS, 10 µL NP-40, 600 µL heptane, 25 µL 16% paraformaldehyde). Ovaries were washed three times in PBST (1x PBS + 0.1% Tween-20), then incubated in blocking solution (PBST + 1% BSA). Then ovaries were incubated in primary antibody diluted in blocking solution at 4° C. Ovaries were then washed three times in PBST and incubated in secondary antibody diluted in blocking solution. After antibody incubation ovaries were washed three times quickly in PBST and mounted with DAPI Fluoromount-G (Thermo Scientific). Antibodies for H3K9me3 (Active Motif, 39161) and C(3)G (Anderson et al. 2005) were used.

### Generation of fluorescence in situ hybridization (FISH) probes

BAC clone (BAC PAC RPCI- 98 library) DNA was extracted using a MIDI-prep kit (Clontech #740410). The probe for the *bw* locus was Clone BACR48M01. BAC DNA was used in nick-translation reaction to create biotinylated probes. Nick translation reaction: 5 µL 10X DNA Pol I buffer, 2.5 µL dNTP mix (1 mM each of dCTP, dATP, dGTP), 2.5 µL biotin-11-dUTP (1 mM), 5.0 µL 100 mM BME, 10 µL of freshly diluted dDNase I, 1 µL DNA Pol I, 1 ug of template DNA, water up to 50 µL. The reaction was incubated in thermocycler at 15° C for four hours. The probe was purified using PCR purification kit (Qiagen) and quantified using Qubit (Thermofisher Q32854), then diluted to 2ng/µL in hybridization buffer (2X saline-sodium citrate (SSC) buffer, 50% formamide, 10% w/v dextran sulfate, 0.8 mg/mL salmon sperm DNA).

AACAC oligonucleotide probe was obtained from Integrative DNA Technologies (IDT, www.idtdna.com). Sequence: Cy3-AACACAACACAACACAACACAACACAACACAACAC.

### Drosophila whole mount ovary IF-FISH

Ovaries were dissected as described above, incubated in fixative buffer for four minutes (100 mM sodium cacodylate (pH7.2), 100 mM sucrose, 40 mM potassium acetate, 10 mM sodium acetate, 10 mM EGTA, 5% paraformaldehyde), washed four times quickly in 2XSSCT (5mL 20X saline sodium citrate (SSC), 50 µL Tween-20, up to 50 mL water), washed 10 minutes in 2X SSCT + 20% formamide, 10 minutes 2X SSCT + 40% formamide, then two times 10 minutes in 2X SSCT + 50% formamide. Ovaries were pre-denatured by incubating at 37° C for 4 hours, 92° C for 3 minutes, 60° C for 20 minutes. Probe(s) was added and ovaries were incubated in a thermocycler at 91° C for three minutes then overnight at 37° C. Ovaries were then washed with 2X SSCT + 50% formamide at 37° C for 1 hour, then in 2X SSCT + 2-% formamide for 10 minutes at room temperature (RT), then in 2X SSCT quickly four times. Ovaries were then incubated in blocking solution (6 mg/mL NGS in 2X SSCT) for four hours, then washed quickly three times in 2X SSCT. Ovaries were incubated overnight in primary antibody diluted in 2X SSCT at room temperature, then washed three times quickly in 2X SSCT, incubated with secondary antibody diluted in 2X SSCT for two hours, then washed three times quickly in 2X SSCT. Ovaries were then incubated with streptavidin (1.5 µL of 488-conjugated streptavidin diluted in 98.5 µL detection solution [0.5 mL 1M Tris, 400 mg BSA, water to 10 mL]) for one hour at room temperature, washed two times quickly in 2X SSCT, one hour in 2X SSCT, then three hours in 2X SSCT. Ovaries were then mounted in DAPI fluoromount. In this work, primary antibody for C(3)G (Anderson et al. 2005) was used.

### Imaging and quantification

Images of whole-mount germaria were taken using a Zeiss LSM710 confocal laser scanning microscope using 40x oil-immersion objective. Images were saved as .czi files and processed using FIJI (ImageJ). Distance between foci for Figure 3 was measured using FIJI. Distances were compared using unpaired t-test.

### Statistical methods

We conducted an analysis of crossover density using a model averaging approach (Burnham, Anderson, and Huyvaert 2011). In this approach, models of varying composition and complexity are weighted according to their ability to fit the data parsimoniously, then averaged to construct predictions and inference. A benefit of this approach is lack of picking one best model when uncertainty exists among a set of candidate models. Similarly, there are no hard *p*-value cutoffs which can be used to artificially exclude weak, but potentially important variables. All statistical analyses were completed using the R language (version 3.6; R Core Team 2019).

The count of crossovers in each chromosome section was modeled with negative binomial regressions fit using maximum likelihood using the MASS library (version 7.3-51.4; Venables and Ripley 2002). All models use a log link function to relate the linear combination of predictor variables to the mean number of crossovers. All models also include an offset variable (a variable whose slope is assumed to be one) of the log(# of number of flies X length of chromosome section). This offset accounts for the different sampling involved in each observation and can be thought of changing the model to one fitting the density of crossovers per fly per section. Prior to fitting, all quantitative variables were centered and standardized by dividing by 2 times the standard deviation of the variable.

The most complex or “global model” included, in addition to the offset, linear additive effects of the density of transposable elements and gene density and a quadratic response to distance from the centromere (distance from the centromere is calculated as distance from the end of the genome assembly for each chromosome arm):

Log(mean # of crossovers) ∼ (distance from centromere + distance from centromere^2^ + transposable element density + gene density)* chromosome identity + log(offset(Fly number * width of chromosome section))

All subsets of this model that included the quadratic effect of distance only when there was a linear effect of distance, were fit. Model selection and averaging were conducted using the MuMIn library (version 1.4.36; Barton 2019). We fit all possible submodels of the global. This led to 150 models being fit. We used the corrected Akaike Information criterion, AICc, as our measure of model performance and selected a final model set based on a 95% confidence set and then calculated model averaged estimates of coefficients and their standard errors. Models which had higher AICc than nested models were excluded based on the recommendation of Richards *et al*. to avoid including overly complex models that do not improve model performance (Richards, Whittingham, and Stephens 2011).

### Data availability

All data necessary for confirming the conclusions in this paper are included in this article and in supplemental figures and tables. *Drosophila* stocks described in this study are available upon request. We have uploaded Supplemental Material to Figshare. Table S1 includes complete data set for crossovers between phenotypic markers in WT and *Blm*. Table S2 includes all primers used for SNP/indel genotyping between isogenized strains of *Drosophila melanogaster*. Table S3 includes all data for mapping of crossovers using the SNP/indel method for WT including chromosome, interval size, number of crossovers, # of genes, # TEs, and total number of flies scored for each interval. Table S4 includes all data for mapping of crossovers using the SNP/indel method for *Blm* mutant. Table S5 includes crossover data between *px* and *sp* for WT and *bw*^*D*^ mutants. Table S6 includes model averaged standardized effect sizes for each chromosome. Table S7 includes the 95% confidence set for wild type chromosome analysis. Table S8 includes modeled average parameters for mutant analysis. Table S9 includes 95% confidence set for *2*L mutant chromosome analysis.

## Results

### Pericentromeric crossover distribution

To gain a deeper understanding of the centromere effect, we sought to more finely map centromere-proximal crossovers. Crossovers near the centromere have classically been mapped using phenotypic markers in the euchromatin on either side of the centromere. Additionally, whole-genome mapping has been used to more precisely map crossovers within the genome (B. A. H. Sturtevant 1915; Miller et al. 2016). However, these methods have caveats that do not allow us to fully understand the distribution of centromere-proximal crossovers. Using phenotypic markers to map crossovers limits resolution to only the most centromere-proximal markers used. Whole-genome mapping provides precise locations of crossovers, but only a handful of centromere-proximal crossovers have been mapped using this method. For example, from whole-genome sequencing of 98 flies only one crossover was mapped between the markers *pr* and *cn* that flank the chromosome *2* centromere (Miller et al. 2016). We therefore develop a method to map a high quantity of crossovers with more precision than phenotypic mapping allows, allowing us to gain a better understanding of the relationship of crossover distribution in euchromatin and the two types of heterochromatin (LR-heterochromatin and HR-heterochromatin).

We collected proximal crossovers between isogenized *Drosophila* chromosomes then more finely mapped these using SNP and indel markers to intervals that range from 0.23 Mb to 1.9 Mb. We mapped approximately 160-300 crossovers per chromosome arm. This mapping shows that crossovers are decreased near the centromere and increase in frequency with distance from the centromere (Figure 1). Interestingly, we see a low frequency of crossovers in the assembled LR-heterochromatin, but crossover frequency goes down to nearly zero in the highly repetitive heterochromatin on every chromosome arm. Of 37,219 total flies scored, only three, all on chromosome *2*, experienced a crossover between the most centromere-proximal SNPs/indels used in our mapping. These crossovers may have occurred within LR-heterochromatin, either proximal to our most proximal markers or in sequences not included in the genome assembly. Alternatively, they may have been within HR-heterochromatin or unique sequences within HR-heterochromatin. We cannot exclude the possibility that these crossovers are mitotic in origin. While we cannot rule out that double crossovers occur within the HR-heterochromatin region, we believe this to be highly unlikely due to the near absence of even single crossovers. The fact that we do see a small amount of crossovers in the less-repetitive heterochromatin was surprising because it has been shown that DSBs do not colocalize with heterochromatic markers (Mehrotra and McKim 2006; see Discussion).

**Figure 1.**
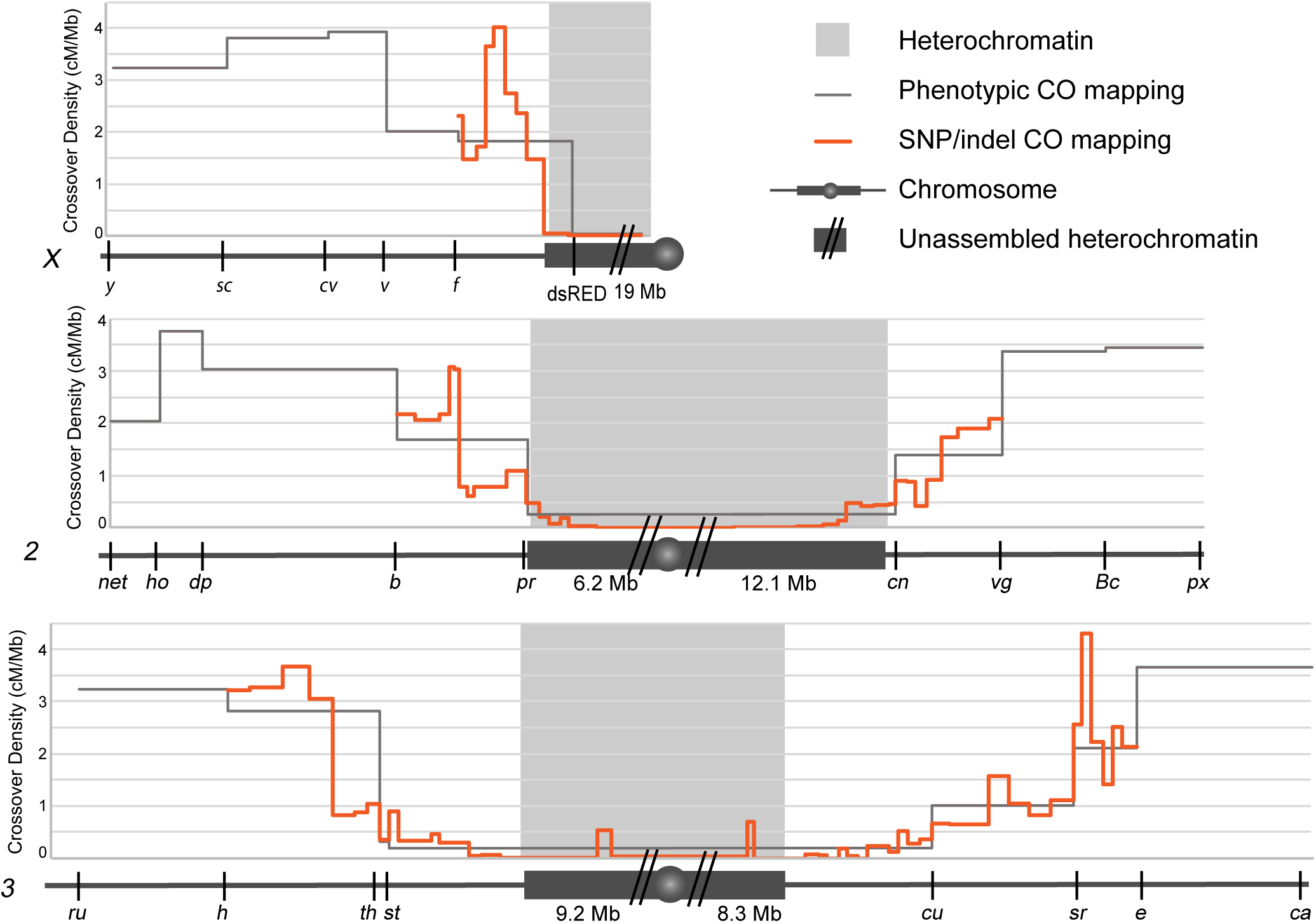
Fine mapping of centromere-proximal crossovers. Chromosomes are represented under each graph (*X, 2, 3*) with euchromatin (dark gray line), heterochromatin (dark gray box), unmapped heterochromatin (dark gray box with two slashes), and the centromere (dark gray circle). Predicted amount of heterochromatin is displayed underneath chromosome for each chromosome arm (values obtained from Hoskins et al. 2002). Heterochromatin boundaries (light gray blocks) are based on H3K9me2 ChIP array boundaries shown in Riddle *et al.* 2011). Phenotypic markers used for mapping crossovers are indicated on each chromosome. Crossover density (cM/Mb) is plotted for crossovers scored between phenotypic markers (gray line) and for crossovers scored using SNP/indel mapping (orange line). Chromosome *X* n=160, Chromosome *2* n=415, Chromosome *3* n=622). For full data set, see Tables S1 and S3.

Fine mapping gives a clearer understanding of crossover distribution near the centromere, but to begin understanding the contribution to this distribution, we explored a mutant that does not experience centromere-proximal crossover suppression.

### A centromere effect mutant separates the centromere effect and the HR-heterochromatin effect

If the centromere effect is genetically controlled, we would anticipate it being possible to identify mutants that do not experience suppression of crossovers near the centromere. Hatkevich *et al.* identified a mutant that they hypothesized does not experience the centromere effect (Hatkevich et al. 2017). *Drosophila* Blm helicase, like *S. cerevisiae* Sgs1, has been proposed to direct DSBs down the meiotic DSB repair pathway to allow the proper crossover patterning (Hatkevich et al. 2017; Zakharyevich et al. 2012; De Muyt et al. 2012). It is hypothesized that *Blm* mutants do not have the centromere effect based on a flat distribution of crossovers and a measure of the strength of the centromere effect (Hatkevich et al. 2017). That study only mapped crossovers using phenotypic markers in the euchromatin on either side of the centromere so we aimed to examine the loss of the centromere effect in *Blm* mutants using the SNP/indel mapping approach. Importantly, *Blm* mutants appear to have normal heterochromatin as evidenced by H3K9me3 staining colocalizing with DAPI dense, heterochromatic regions (Figure 2A).

**Figure 2.**
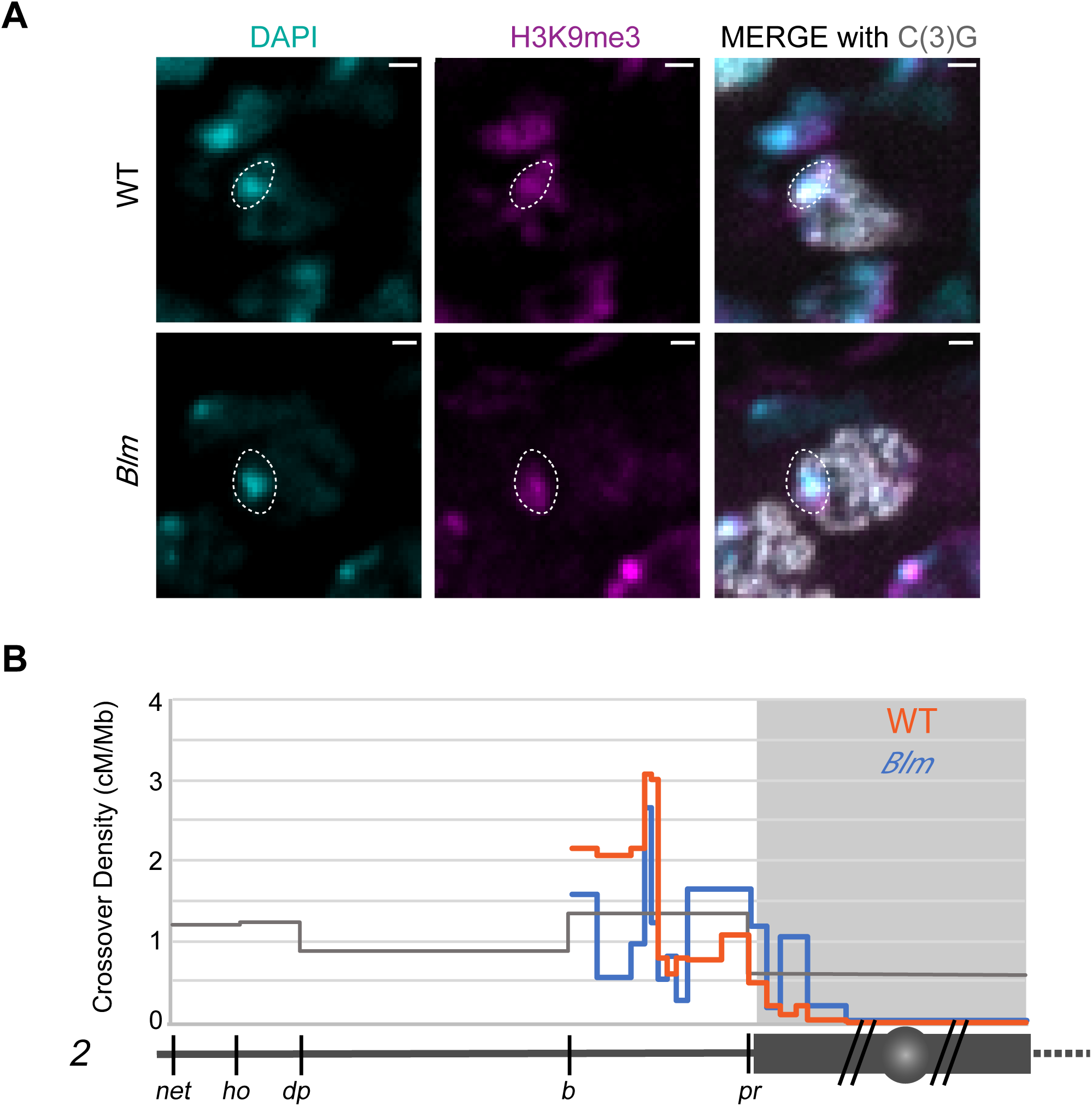
Fine mapping of centromere-proximal crossovers in *Blm* mutants. (A) Heterochromatin staining in *Blm* mutants and WT. DAPI staining for DNA is shown in the left panels, H3K9me3 staining for heterochromatin shown in the middle panels, and the right panels are merged images. The dotted circle outlines a DAPI region that overlaps with heterochromatin showing that *Blm* mutants have normal localization of heterochromatin. (B) SNP/indel mapping as shown in Figure 1 for chromosome *2* with euchromatin (dark gray line), heterochromatin (dark gray box), unmapped heterochromatin (dark gray box with two slashes), and the centromere (dark gray circle). Phenotypic markers are depicted under the chromosome. Plotted on the graph is crossover density (cM/Mb) for phenotypic markers (gray), WT SNP/indel mapping (orange), and *Blm* SNP/indel mapping (blue). Heterochromatin shown by light gray box. For full data sets, see Tables S1 and S4.

SNP/indel mapping of *Blm* mutants reveals a relatively flat distribution of crossovers throughout the chromosome arm and into the assembled heterochromatin (Figure 2B). *Blm* mutants experience no crossovers within the HR-heterochromatin, as in wildtype. From these results, we hypothesize that the suppression of crossovers can be separated into two phenomena: the HR-heterochromatin effect, defined as the virtual absence of crossovers within highly-repetitive heterochromatin, and the centromere effect, which has a dissipating effect with distance from the centromere. We hypothesize that the HR-heterochromatin effect is likely due to the absence of DSBs in this region, whereas the centromere effect is likely a regulation of DSB repair outcome.

### Heterochromatin alone does not produce a centromere effect

We sought to test whether the HR-heterochromatin effect and centromere effect can be separated by measuring recombination around a heterochromatic locus that is not located near the centromere. We do this by using the *bw*^*D*^ mutation, which has an insertion of about 2Mb of heterochromatin in the *bw* locus on distal chromosome *2*R (Slatis 1955; Dernburg et al. 1996) (Figure 3A). This mutation causes dominant suppression of the *bw* gene by pairing with its homolog and causing localization near the pericentromeric heterochromatin of chromosome *2* (Dreesen, Henikoff, and Loughney 1991; S Henikoff and Dreesen 1989; Dernburg et al. 1996; Steven Henikoff, Jackson, and Talbert 1995). We used this tool to answer two questions: First, does an insertion of heterochromatin located far from the centromere suppress crossovers in adjacent intervals? Second, does spatial proximity to pericentromeric heterochromatin within the nucleus suppress crossovers?

**Figure 3.**
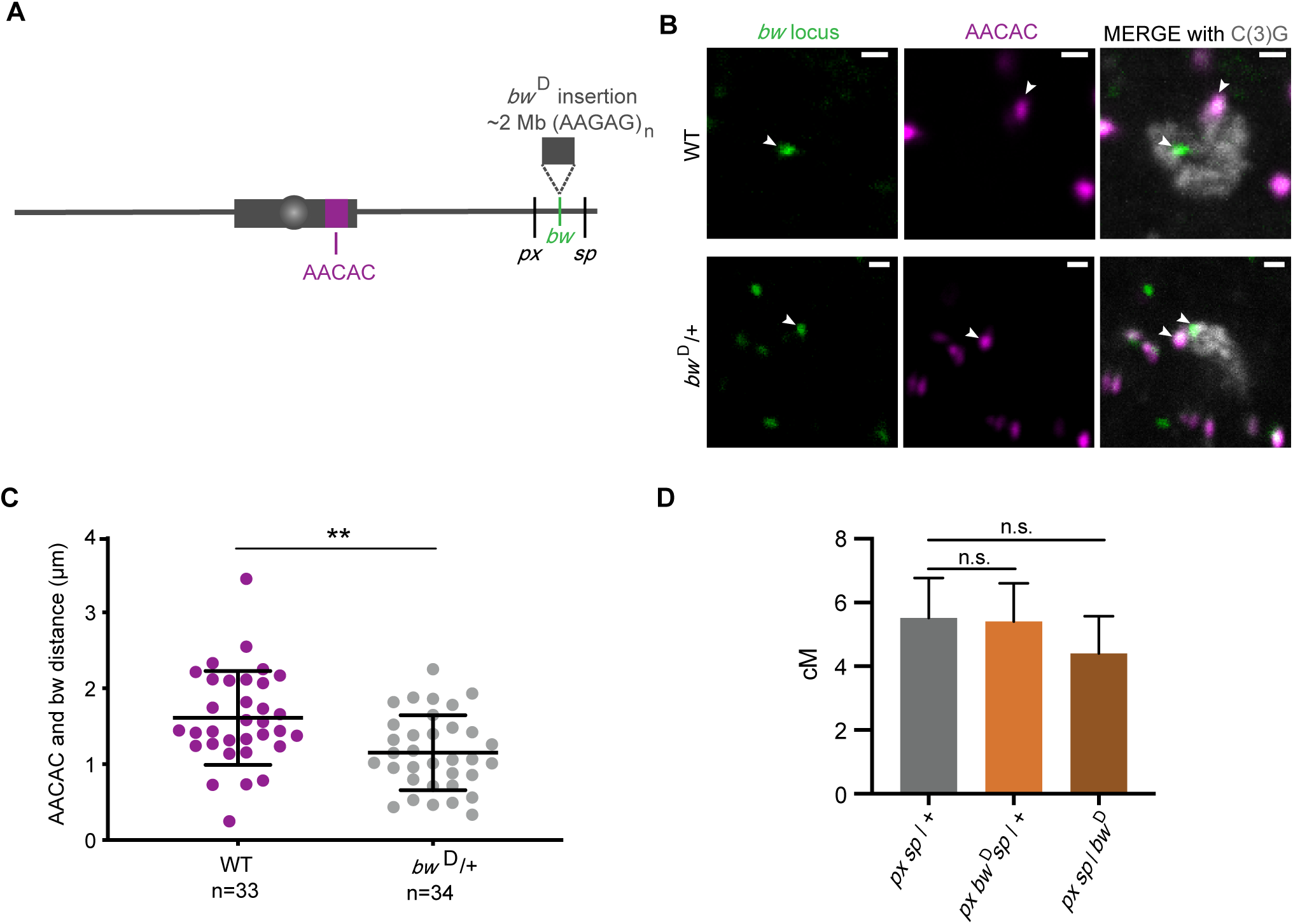
Insertion of a block of heterochromatin does not decrease crossovers. (A) Schematic of the *bw*^*D*^ mutation and AACAC locus used for staining. (B) Representative staining for *bw* locus (left panels), AACAC locus (middle panels), and C(3)G to identify meiotic cells (merged with foci in the right panels). White arrows point to the foci in all images. (C) Quantification of the distance between foci in WT and *bw*^*D*^/+. (** *p* < 0.001). (D) Recombination between *px* and *sp* represented in cM for *px sp /* + (5.52 cM n= 1287), *px bw*^*D*^ *sp /* + (5.4 cM n= 1363), *px sp / bw*^*D*^ (4.4 cM n= 1197). Error bars represent 95% confidence intervals. (*px sp /* + versus *px bw*^*D*^ *sp /* +, n.s. *p* = 0.86) (*px sp /* + versus *px sp / bw*^*D*^, n.s. *p*=0.32). For full data set, see Table S5.

We first asked whether the heterochromatic insertion of *bw*^*D*^ causes nuclear localization of the locus near clustered pericentromeric heterochromatin in meiotic cells in the same fashion as it does in somatic cells. We used a probe for the *bw* locus and a probe for a repeat in the pericentromeric heterochromatin of chromosome *2* (AACAC), as well as a marker of meiotic cells, C(3)G, a component of the synaptonemal complex (Figure 3B). We then measured the distance between the two foci in meiotic cells and see that the distance between the *bw* locus and AACAC heterochromatin locus is significantly shorter in *bw*^*D*^ compared to WT (*p* < 0.001) (Figure 3C). This suggests that the heterochromatic insertion in *bw*^*D*^ does localize near the pericentromeric heterochromatin in meiotic cells.

We measured recombination between phenotypic markers on either side of the *bw* locus, *px* and *sp*. The *px* gene is located at 2R:22.5 Mb; *sp* is not mapped to the genome but is between *or* at 2R:24.0 Mb and *Kr* at 2R:25.2 Mb, so the distance between *px* and *sp* is between 1.5-2.7 Mb (Thurmond *et al.* 2019). If the heterochromatic insertion in *bw*^*D*^ leads to suppression of crossovers in adjacent regions we would expect to see a decrease in crossovers between *px* and *sp*. We assume there are no crossovers within the heterochromatin of the *bw*^*D*^ mutation since we measured crossovers in *bw*^*D*^ heterozygous background. Surprisingly, there was not a significant difference in number of crossovers with the *bw*^*D*^ insertion being either *cis* or *trans* to *px* and *sp* (*p* = 0.86 and *p* = 0.32) (Figure 3D), suggesting that the heterochromatin insertion does not cause a decrease in crossovers in the adjacent regions and that spatial proximity to the pericentromeric heterochromatin compartment of the nuclease does not have a strong effect on crossing over.

### Examination of contributions to the centromere effect

The results with *bw*^*D*^ suggest that the centromere effect is not due solely to proximity to pericentromeric heterochromatin, so we asked whether other genomic features contribute to the centromere effect. Transposable element (TE) density and gene density have been suggested to influence crossover rates genome-wide in other organisms (Bartolome, Maside, and Charlesworth 2001; Kent, Uzunović, and Wright 2017; Bartolome and Maside 2004). TEs are middle-repetitive elements found throughout the genome but are most abundant within LR-heterochromatin adjacent to euchromatin (Carmena and González 1995; M. T. Yamamoto et al. 1990). Conversely, genes are less abundant in the LR-heterochromatin than in the euchromatin. In *Arabidopsis thaliana* crossovers are negatively correlated with TE density and positively correlated with gene density (Giraut et al. 2011). Therefore, we searched for correlations between crossover distribution and distance from the centromere, TE density and gene density.

Figures 4A and B show TE and gene density overlaid with our SNP/indel mapping of proximal crossovers. We modeled how distance from the centromere, TE density, and gene density contribute to the variation seen in crossover distribution (Figure 4C). Two models were selected in the 95% confidence set (Tables S6 and S7). All predictor variables were included in this final set indicating statistically important effects of distance from the centromere, TE density and gene density that varied across chromosomes. Unless otherwise stated, all effects mentioned have 95% confidence intervals that do not overlap zero. For all chromosomes except *X*, distance from the centromere had a positive effect and a negative squared distance term. Two chromosome arms, *2R* and *X*, had positive effects of gene density; on *3R*, a negative effect was found with 95% confidence intervals just overlapping zero, suggesting a potential negative effect. In general, standardized effect sizes for gene density were lower than for distance from the centromere. For TE density all chromosomes but *X* had 95% confidence intervals that did not overlap zero. The effect was dramatically negative in *3R* with a negative standardize effect size of magnitude over three times greater than the next effect size. Other chromosomes had smaller magnitude effect size, being negative for *2L, 3L*, and *3R*, but positive for *2R*. This modeling shows that TE and gene density do decrease the variation seen in the model; however, they do not fully explain the model produced and there is leftover effect of distance from the centromere. These results support the idea that centromere-proximal crossover distribution is dictated not only by genomic features such as TE or gene density, but that there is some factor suppressing crossover rate that decreases with distance from the centromere.

**Figure 4.**
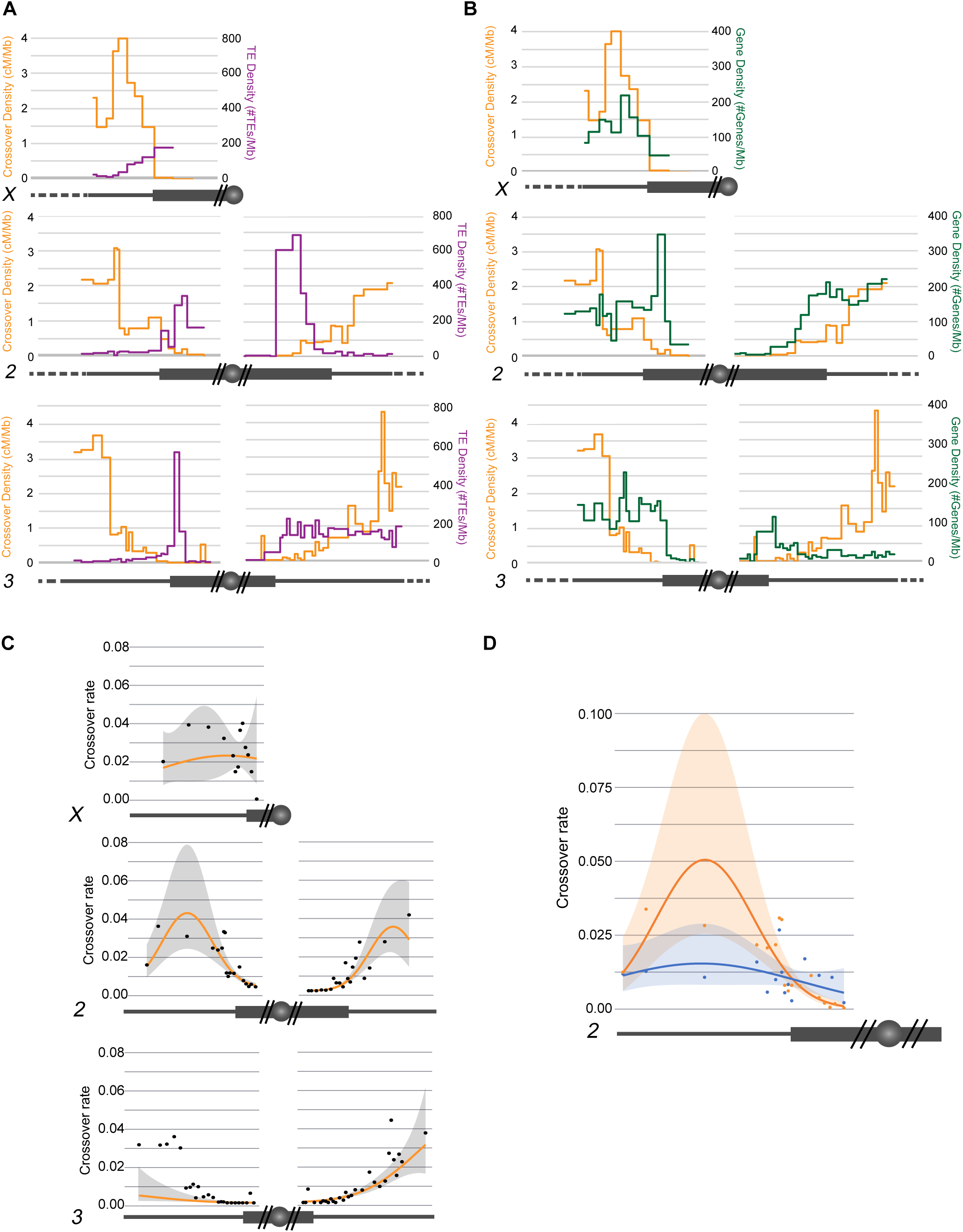
Distribution of TE and gene density. Chromosomes are depicted under each graph (*X, 2, 3*) with euchromatin (dark gray line), heterochromatin (dark gray box), unmapped heterochromatin (dark gray box with two slashes), and the centromere (dark gray circle). (A and B) Crossover distribution from SNP/indel mapping (orange) represented on left axis as cM/Mb. (A) Transposable element (TE) density (purple) plotted on right axis as number of TEs/Mb. (B) Gene density (green) plotted on right axis as number of genes/Mb. (C) Crossover rate in relation to distance from the centromere. Observed data are plotted along with modeled marginal relationship with distance from the centromere. For the marginal predictions, gene density and transposable element density were set at their mean value across each chromosome. (D) Crossover rate in relation to distance from the centromere for chromosome *2*L for *Blm* mutant and WT. Observed data are plotted along with modeled marginal relationship with distance from the centromere. For the marginal predictions, gene density and transposable element density were set at their mean value across each chromosome. For statistical analyses, see Tables S6-S9.

We applied the same modeling methods to the *Blm* mutant to understand if *Blm* mutants truly do not have a centromere effect and to what extent TE and gene density play a role in crossover distribution in *Blm* mutants (Figure 4D). Two models were selected in the 95% confidence set (Tables S8 and S9). There was no effect of gene density in either wild type or mutant, consistent with analysis of the wild-type chromosomes. In the wild type, all remaining modeled effects (distance, distance^2^ and TE density) had 95% confidence intervals that did not overlap zero. In the mutant, no effect size had confidence intervals that didn’t overlap zero, suggesting that none of them were valuable predictors of crossover rate. While we cannot prove zero effect, the best estimated effect of distance in the mutant is less than one quarter that of the wild-type (Table S8). These results support the hypothesis that *Blm* mutants experience a much weaker centromere effect, if any, and that the crossover distribution in *Blm* is not demonstrably under the influence of distance from the centromere or chromosome characteristics. Importantly, these results provide more evidence that centromere-proximal crossover suppression is mediated both by the HR-heterochromatin effect and an effect whose strength varies with distance to the centromere.

## Discussion

### Two Contributions to Suppression of Proximal Crossovers

Our mapping of a large number of proximal crossovers in both wild-type flies and *Blm* mutants leads us to propose a model for centromere-proximal crossover suppression (Figure 5). In this model crossovers are completely suppressed in HR-heterochromatin due to the absence of DSBs. Adjacent to this region the centromere effect strongly suppresses crossovers, but that suppression dissipates with distance from the centromere until a region in the euchromatin where crossovers rise steeply to peak around the middle of each chromosome arm (orange line). In the *Blm* mutant (blue line), the HR-heterochromatin effect is still intact but the centromere effect is lost: crossover density is relatively even throughout the assembled LR-heterochromatin and euchromatin. We conclude that pericentromeric crossover suppression is achieved by both HR-heterochromatin suppression and a centromere effect, and these two processes are separable.

**Figure 5.**
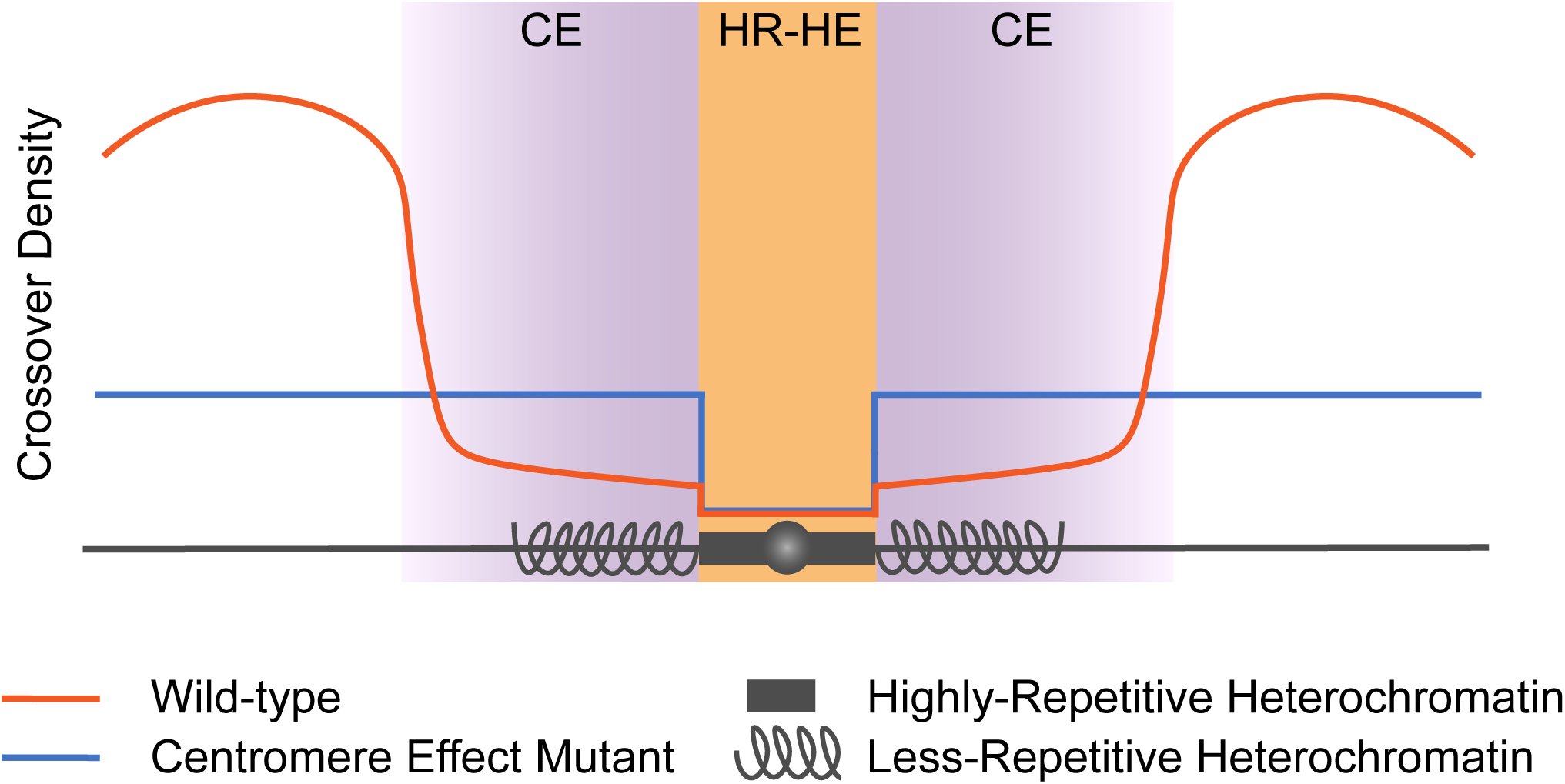
Model of the Centromere Effect and HR-Heterochromatin effect. The HR-Heterochromatin Effect (HR-HE) (orange) and Centromere Effect (CE) (purple) are both responsible for suppressing crossovers in the pericentromeric region. The HR-HE completely suppresses crossovers in the highly-repetitive heterochromatin (dark gray box), but does not have an effect outside of this region. The CE starts in the less-repetitive heterochromatin (spiral gray lines) and extends into the euchromatin of each arm (straight gray lines). Representative crossover distribution is shown for WT (orange) and a centromere effect mutant (blue).

### Heterochromatin effect suppresses crossovers

Heterochromatin has long been thought to contribute to centromere-proximal suppression of crossovers, but the specifics of where this suppression occurred were unknown until now. In this study, we used a centromere effect mutant (*Blm*) that still has normal heterochromatic marks to show that the heterochromatin effect impacts the highly-repetitive heterochromatin but not the adjacent less-repetitive heterochromatin. This was a surprising result because a previous cytological study found that a marker of DSBs never colocalized with a heterochromatin marker (Mehrotra and McKim 2006). It is possible that, although DSBs do occur within less-repetitive heterochromatin, they are at a lower density than in euchromatin (perhaps by being excluded from TE sequences) so the sample size in the previous study was insufficient to detect these relatively rare events.

We used *bw*^*D*^ to test whether HR-heterochromatin distant from the centromere exerts a centromere effect. The lack of an effect on crossing over between *px* and *sp* suggests that HR-heterochromatin is not sufficient to reduce crossovers in flaking regions. We could not assay the effects of homozygosity for *bw*^*D*^ because homozygotes were inviable, even for chromosomes with the closely-spaced markers *px* and *sp* recombined onto *bw*^*D*^. Slatis (1955) conducted similar experiments and reported a decrease in crossovers in flies heterozygous and homozygous for *bw*^*D*^. This may suggest that HR-heterochromatin does affect adjacent euchromatin even when distant from the centromere. However, Slatis also reported a decrease in *bw*^*D*^ heterozygotes, in contrast to our findings; the reasons for this difference are unknown. After the genomic location of *sp* is determined, it would informative to revisit these studies and map crossovers between *px, bw*^*D*^, and *sp* more precisely.

Why can crossovers occur within the less-repetitive heterochromatin, but not the highly-repetitive heterochromatin? One reason could be differential access of DSB machinery to the DNA. Perhaps HR-heterochromatin is more densely packed than LR-heterochromatin and does not allow access of the DSB machinery. Additionally, there could be different heterochromatic marks or protein machinery in these regions that differentially regulate DSBs or crossover formation. Future studies could be aimed at determining functional differences between LR- and HR-heterochromatin.

In the centromere effect mutant, *Blm*, we show that crossovers do not occur within the highly-repetitive heterochromatin, but they do occur outside of that boundary at a higher frequency than wildtype crossovers. This suggests that DSBs are still occurring within the less-repetitive heterochromatin at a rate similar to the euchromatin, but that in wildtype, they are more frequently being converted to noncrossovers instead of crossovers. However, Westphal and Reuter reported an increase in centromere-proximal crossovers in *Su(var)* mutants, which presumably cause heterochromatin to assume a more open structure (Westphal and Reuter 2002). This result suggests that the closed structure of heterochromatin can suppress crossovers, but that is in opposition to the result we see with the centromere effect mutant that still has normal heterochromatic marks, but allows more crossovers within the heterochromatic region. There are two possible explanations that could explain these opposing results. The first is that the *Blm* mutation is altering heterochromatin structure in a way that we did not detect cytologically. In this case, it would be interesting to look at distribution of heterochromatin marks in meiotic cells of the mutant, but this is currently not feasible because we do not have a way of isolating meiotic cells for studies such as ChIP analysis of heterochromatic marks. Additionally, in the *Su(var)* mutants, perhaps the opening of heterochromatin in both the less- and highly-repetitive heterochromatin allows crossovers to occur within the highly-repetitive heterochromatin. This could show us separation of the centromere effect and HR-heterochromatin effect by retaining the centromere effect but disrupting the HR-heterochromatin effect. It would be informative to conduct our SNP/indel mapping on crossovers in a *Su(var)* mutant.

*Blm* mutants also experience crossovers on chromosome *4*, which normally never has crossovers (Hatkevich et al. 2017). We hypothesized that chromosome *4* does not have crossovers because of a very strong centromere effect, which is lost in *Blm* mutants (Hartmann and Sekelsky 2017); the results reported here support this hypothesis. It would be interesting to finely map crossovers on chromosome *4* in *Blm* mutants to determine if there is a flat distribution and see if there is a separable HR-heterochromatin effect on this chromosome as well (this would require a marker to the left of the centromere).

### Recombination and genomic features

The relationship between gene density, TE density, and recombination rate has been a long-standing discussion (reviewed in Kent *et al.* 2017). It is difficult to parse out these relationships because there are many factors influencing distribution of TEs, genes, and crossovers. It has been argued that the distribution of TEs and genes is in part dictated by recombination. For example, higher recombination could be favored in regions of high gene density to promote greater genetic diversity within populations. Conversely, lower recombination rates in regions of high TE density could help to prevent ectopic recombination between similar TE sequences in different genomic locations. The high density of TEs in proximal or heterochromatic regions could actually result from the low recombination rate preventing removal of TEs (Bartolome and Maside 2004). Recombination might also be directly silenced within TE sequences. Miller *et al.* (Miller et al. 2016) reported that crossovers can occur within TEs, but less frequently than would be expected. It has been suggested that active silencing of TEs could lead to the silencing or suppression of recombination around those regions (Kent, Uzunović, and Wright 2017). Therefore, it is difficult to determine whether or how TE density and gene density affect recombination rates. Our data support results seen previously in that TE density is increased in areas of low recombination and gene density is increased in areas of high recombination. When we factor these variables into models of crossover distribution, we see a strong impact of TE density on crossover rate. One caveat of our studies is that transposable elements have been shown to vary between different strains of *Drosophila* and we have based these analyses off the transposable element distribution within the *Drosophila melanogaster* reference sequence (Ananiev et al. 1984; Rahman et al. 2015). With advances in long-read sequencing technology, it might be possible in the future to do studies similar to ours but in strains in which LR-heterochromatin has been assembled *de novo*.

## Conclusion

In conclusion, we find that centromere-proximal crossover suppression is a result of two separable mechanisms: an HR-heterochromatin effect that suppresses crossovers in highly-repetitive pericentromeric heterochromatin, and the centromere effect that suppresses proximal crossovers in a manner that dissipates with increasing distance from the centromere. The HR-heterochromatin effect is likely due to the absence of DSBs with satellite sequences, presumably a direct consequence of chromatin structure. In contrast, the mechanism of the centromere effect is unknown. This work is the first in-depth examination of the centromere effect since it was first described, and these findings provide the groundwork for future mechanistic studies of the centromere effect.

## Acknowledgements

We thank Juan Carvajal Garcia for comments on the manuscript. This work was supported by a grant from the National Institute of General Medical Sciences to J.S. under award 1R35 GM-118127.

## Author Summary

Crossovers are essential for the proper segregation of chromosomes, as is their accurate positioning along the chromosome. Crossovers are normally reduced near the centromere, and this suppression has been referred to as the centromere effect. However, very little is known about the centromere effect and there has been no insight into the mechanism. Here, we investigate mechanisms behind centromere-proximal crossover suppression, and show that this suppression is mediated by both the highly-repetitive heterochromatin effect that completely suppresses crossovers within highly-repetitive heterochromatin and the centromere effect, which suppresses crossovers with a dissipating effect with distance from the centromere.

## References

Ananiev, E. V., V. E. Barsky, Yu V. Ilyin, and M. V. Ryzic. 1984. The arrangement of transposable elements in the polytene chromosomes of *Drosophila melanogaster*. Chromosoma 90 (5): 366–77. https://doi.org/10.1007/BF00294163.

Anderson, L. K., S. M. Royer, S. L. Page, K. S. McKim, A. Lai, M. A. Lilly, and R. S. Hawley. 2005. Juxtaposition of C(2)M and the transverse filament protein C(3)G within the central region of *Drosophila* synaptonemal complex. Proceedings of the National Academy of Sciences 102 (12): 4482–87. https://doi.org/10.1073/pnas.0500172102.

Ashburner, M. 1980. Some Aspects of the Structure and Function of the Polytene Chromosomes of the Diptera. Insect Cytogenetics 10: 65–84.

Bartolome, Carolina, and Xulio Maside. 2004. The lack of recombination drives the fixation of transposable elements on the fourth chromosome of *Drosophila melanogaster*. Genet Res 83 (2): 91–100. https://doi.org/10.1017/S0016672304006755.

Bartolome, Carolina, Xulio Maside, and Brian Charlesworth. 2001. On the abundance and distribution of transposable elements in the genome of *Drosophila melanogaster*. Mol. Biol. Evol. 19: 926–37.

Barton, K. 2019. MuMIn: Multi-Model Inference. https://cran.r-project.org/package=MuMIn.

Beadle, G. W. 1932. A possible influence of the spindle fibre on crossing-over in *Drosophila*. Genetics 18: 160–65.

Berchowitz, Luke E, and Gregory P Copenhaver. 2010. Genetic interference: Don’t stand so close to me. Current Genomics 11 (2): 91–102. https://doi.org/10.2174/138920210790886835.

Burnham, Kenneth P., David R. Anderson, and Kathryn P. Huyvaert. 2011. AIC model selection and multimodel inference in behavioral ecology: Some background, observations, and comparisons. Behavioral Ecology and Sociobiology 65 (1): 23–35. https://doi.org/10.1007/s00265-010-1029-6.

Bushnell, B. 2014. BBMap. 2014. https://sourceforge.net/projects/bbmap/.

Carmena, Mar, and Cayetano González. 1995. Transposable elements map in a conserved pattern of distribution extending from beta-heterochromatin to centromeres in *Drosophila melanogaster*. Chromosoma 103 (10): 676–84. https://doi.org/10.1007/BF00344228.

Danecek, Petr, Adam Auton, Goncalo Abecasis, Cornelis A. Albers, Eric Banks, Mark A. DePristo, Robert E. Handsaker, et al. 2011. The variant call format and VCFtools. Bioinformatics 27 (15): 2156–58.

Dernburg, Abby F., Karl W. Broman, Jennifer C. Fung, Wallace F. Marshall, Jennifer Philips, David A. Agard, and John W. Sedat. 1996. Perturbation of nuclear architecture by longdistance chromosome interactions. Cell 85 (5): 745–59.

Dreesen, T D, S Henikoff, and K Loughney. 1991. A pairing-sensitive element that mediates trans-inactivation is associated with the *Drosophila* Brown Gene. Genes and Development 5: 331–40.

Filion, Guillaume J., Joke G. van Bemmel, Ulrich Braunschweig, Wendy Talhout, Jop Kind, Lucas D. Ward, Wim Brugman, et al. 2010. Systematic protein location mapping reveals five principal chromatin types in *Drosophila* cells. Cell 143 (2): 212–24.

Gall, Joseph G. 1973. Repetitive DNA in *Drosophila*. Molecular Cytogenetics, 59–60.

Giraut, Laurène, Matthieu Falque, Jan Drouaud, Lucie Pereira, Olivier C. Martin, and Christine Mézard. 2011. Genome-wide crossover distribution in *arabidopsis thaliana* meiosis reveals sex-specific patterns along chromosomes. PLoS Genetics 7 (11).

Hartmann, Michaelyn A, and Jeff Sekelsky. 2017. The absence of crossovers on chromosome 4 in drosophila melanogaster?: Imperfection or interesting exception? Fly 6934 (May): 4–11.

Hatkevich, Talia, Kathryn P Kohl, Susan Mcmahan, Michaelyn A Hartmann, Andrew M Williams, Jeff Sekelsky, Talia Hatkevich, et al. 2017. Bloom syndrome helicase promotes meiotic crossover patterning and homolog disjunction. Current Biology 27: 96–102.

Henikoff, S, and T D Dreesen. 1989. Trans-Inactivation of the Drosophila Brown Gene: Evidence for transcriptional repression and somatic pairing dependence. Proceedings of the National Academy of Sciences of the USA 86 (17): 6704–8. https://doi.org/10.1073/pnas.86.17.6704.

Henikoff, Steven, Jeffrey M Jackson, and Paul B Talbert. 1995. Distance and pairing effects on the brown dominant heterochromatic element in *Drosophila*. Genetics 140: 1007–17.

Hoskins, Roger A., Joseph W. Carlson, Kenneth H. Wan, Soo Park, Ivonne Mendez, Samuel E. Galle, Benjamin W. Booth, et al. 2015. The Release 6 reference sequence of the *Drosophila melanogaster* genome. Genome Research 25 (3): 445–58. https://doi.org/10.1101/gr.185579.114.

Hoskins, Roger A., Christopher D Smith, Joseph W Carlson, a Bernardo Carvalho, Aaron Halpern, Joshua S Kaminker, Cameron Kennedy, et al. 2002. Heterochromatic sequences in a *Drosophila* whole-genome shotgun assembly. Genome Biology 3.

Kent, Tyler V., Jasmina Uzunović, and Stephen I. Wright. 2017. Coevolution between Transposable elements and recombination. Philosophical Transactions of the Royal Society B: Biological Sciences 372 (1736).

Khost, DE, DG Eickbush, and AM Larracuente. 2017. Single molecule long read sequencing resolves the detailed structure of complex satellite DNA loci in *Drosophila melanogaster*. Genome Research 27: 709–21.

Koehler, K E, C L Boulton, H E Collins, R L French, K C Herman, S M Lacefield, L D Madden, C D Schuetz, and R S Hawley. 1996. Spontaneous X chromosome MI and MII nondisjunction events in *Drosophila melanogaster* oocytes have different recombinational histories. Nature Genetics 14 (4): 406–14.

Koehler, K E, R S Hawley, S Sherman, and T Hassold. 1996. Recombination and nondisjunction in humans and flies. Human Molecular Genetics 5 Spec No (8): 1495–1504.

Kohl, Kathryn P., Corbin D. Jones, and Jeff Sekelsky. 2012. Evolution of an MCM complex in flies that promotes meiotic crossovers by blocking BLM helicase. Science 338.

Laird, C., M Hammond, and M. Lamb. 1987. Polytene chromosomes of *Drosophila*. In Chromosomes Today 9: 40–47.

Lake, Cathleen M., and R. Scott Hawley. 2016. Becoming a crossover-competent DSB. Seminars in Cell and Developmental Biology 54: 117–25.

Lamb, N E, S B Freeman, a Savage-Austin, D Pettay, L Taft, J Hersey, Y Gu, et al. 1996. susceptible chiasmate configurations of chromosome 21 predispose to non-disjunction in both maternal meiosis I and meiosis II. Nature Genetics 14: 400–405.

Li, Heng. 2011. A statistical framework for snp calling, mutation discovery, association mapping and population genetical parameter estimation from sequencing data. Bioinformatics 27 (21): 2987–93.

Li, Heng, Bob Handsaker, Alec Wysoker, Tim Fennell, Jue Ruan, Nils Homer, Gabor Marth, Goncalo Abecasis, and Richard Durbin. 2009. The sequence alignment/map format and SAMtools. Bioinformatics 25 (16): 2078–79.

Mather, K. 1937. The determination of position in crossing-over. II. The chromosome lengthchiasma frequency relation. Cytologia, 514–26.

Mather, K. 1939. Crossing over and heterochromatin in the X chromosome of *Drosophila melanogaster*. Genetics 24 (May): 413–35.

McVey, Mitch, Sabrina L. Andersen, Yuri Broze, and Jeff Sekelsky. 2007. Multiple functions of Drosophila BLM helicase in maintenance of genome stability. Genetics 176 (4): 1979–92..

Mehrotra, S., and K. S. McKim. 2006. Temporal analysis of meiotic dna double-strand break formation and repair in Drosophila females. PLoS Genetics 2 (11): 1883–97.

Miklos, George L. Gabor, and J.N. Cotsell. 1990. Chromosome structure at interfaces between major chromatin types: alpha- and beta-heterochromatin. BioEssays 12 (1): 1–6.

Miller, Danny E, Clarissa B Smith, Nazanin Yeganeh Kazemi, Alexandria J Cockrell, and Alexandra V Arvanitakas. 2016. Whole-genome analysis of individual meiotic events in *Drosophila melanogaster* reveals that noncrossover gene conversions are insensitive to interference and the centromere effect. Genetics 203 (1): 159–71.

Muyt, Arnaud De, Lea Jessop, Elizabeth Kolar, Anuradha Sourirajan, Jianhong Chen, Yaron Dayani, and Michael Lichten. 2012. BLM helicase ortholog Sgs1 is a central regulator of meiotic recombination intermediate metabolism. Molecular Cell 46 (1): 43–53.

R Core Team. 2019. R: A Language and environment for statistical computing. r foundation for statistical computing, Vienna, Austria. 2019. https://www.r-project.org/.

Rahman, Reazur, Gung Wei Chirn, Abhay Kanodia, Yuliya A. Sytnikova, Björn Brembs, Casey M. Bergman, and Nelson C. Lau. 2015. Unique Transposon Landscapes Are Pervasive across Drosophila Melanogaster Genomes. Nucleic Acids Research 43 (22): 10655–72.

Richards, Shane A., Mark J. Whittingham, and Philip A. Stephens. 2011. Model selection and model averaging in behavioural ecology: The utility of the IT-AIC framework. Behavioral Ecology and Sociobiology 65 (1): 77–89.

Riddle, Nicole C., Aki Minoda, Peter V. Kharchenko, Artyom A. Alekseyenko, Yuri B. Schwartz, Michael Y. Tolstorukov, Andrey A. Gorchakov, et al. 2011. Plasticity in patterns of histone modifications and chromosomal proteins in *Drosophila* heterochromatin. Genome Research 21 (2): 147–63.

Slatis, H M. 1955. A reconsideration of the brown-dominant position effect. Genetics 40: 246–51.

Sturtevant, A H. 1913. A third group of linked genes in Drosophila ampelophila. Science 37 (965): 990–92.

Sturtevant, A H, and G W Beadle. 1936. The relations of inversions in the X chromosome of *Drosophila melanogaster* to crossing over and disjunction. Genetics 21: 554–604.

Sturtevant, By A H. 1915. The behavior of the chromosomes as studied through linkage. Zeitschrift Für Induktive Abstammungs- Und Vererbungslehre 13: 234–87.

Thurmond, J, JL Goodman, VB Strelets, H Attrill, LS Gramates, SJ Marygold, BB Matthews, et al. 2019. FlyBase 2.0: The Next Generation. Nucleic Acids Res. 47 (D1): D759–65.

Venables, William N., and Brian D. Ripley. 2002. *Modern Applied Statistics with S*. Modern Applied Statistics with S. Fourth ed. Springer.

Wang, Shunxin, Denise Zickler, Nancy Kleckner, and Liangran Zhang. 2015. Meiotic crossover patterns: obligatory crossover, interference and homeostasis in a single process. Cell Cycle 14 (3): 305–14.

Westphal, Thomas, and Gunter Reuter. 2002. Recombinogenic effects of suppressors of position-effect variegation in *Drosophila*. Genetics 160 (2): 609–21.

Yamamoto, M. T., A. Mitchelson, M. Tudor, K. O’Hare, J. A. Davies, and G. L. Gabor Miklos. 1990. Molecular and cytogenetic analysis of the heterochromatin-euchromatin junction region of the *Drosophila melanogaster X* chromosome using cloned DNA sequences. Genetics 125 (4): 821–32.

Yamamoto, Masatoshi, and George L. Gabor Miklos. 1978. Genetic Studies on Heterochromatin in *Drosophila melanogaster* and Their Implications for the Functions of Satellite DNA. Chromosoma 98: 71–98.

Zakharyevich, Kseniya, Shangming Tang, Yunmei Ma, and Neil Hunter. 2012. Delineation of joint molecule resolution pathways in meiosis identifies a crossover-specific resolvase. Cell 149 (2): 334–47.

